## Supplementary material for "NANOG assembles into self-limiting aging micelles that drive a sol-gel transition and modulate DNA dynamics": si text

### I. THE MPIPI MODEL

To simulate NANOG molecules, we employed the Mpipi coarse-grained model [1]. In Mpipi, each amino acid is represented as a single isotropic bead characterized by its mass, diameter and charge. The total potential energy is expressed as

$$V_{\rm Mpipi} = \sum_{\rm bonds} V_{\rm bond} + \sum_{\rm pairs} (V_{\rm elec} + V_{\rm pair}), \qquad \text{(S1)}$$

where $V_{\rm bond}$ is a harmonic term enforcing chain connectivity, $V_{\rm elec}$ is a screened Coulomb potential described by the Debye–Hückel formalism, and $V_{\rm pair}$ is a Wang–Frenkel potential that governs non-bonded residue interactions.

Model parameters were derived from a combination of all-atom potential of mean force (PMF) calculations at 150 mM salt concentration and sequence statistics from bioinformatics datasets. This hybrid parameterization enables Mpipi to capture key residue-specific noncovalent interactions, including hydrophobic, cation–($\pi$), and ($\pi$)–($\pi$) interactions. Notably, the model distinguishes between arginine and lysine residues, reflecting their distinct propensities for electrostatic and aromatic interactions.

Mpipi has been validated against a range of experimental observables, including protein radii of gyration and phase diagrams, and has demonstrated robust predictive accuracy for both protein–RNA and protein–protein phase separation.

### II. DETAILS OF NANOG COARSE-GRAINED SIMULATIONS

The mouse NANOG protein comprises $N = 305$ amino acids (sequence shown in Table S1) and contains three major domains: the N-terminal domain (ND), the homeodomain (HD), and the C-terminal domain (CD). The CD itself is subdivided into CD1, the tryptophan repeat (WR), and CD2 [2] (see Fig. **1**(a) in the main text). We note that, based on sequence alone, NANOG is expected to be weakly negatively charged. Additionally, the WR region contains no charged amino acids. Thus, the original parameterization [1] of the Mpipi model is expected to provide a reliable description of the system.

#### Computation of radius of gyration in dilute conditions

Coarse-grained molecular dynamics simulations were performed for a single polymer of either wild-type NANOG or the W10A mutant under dilute conditions. Each molecule was placed in a cubic simulation box of side length $L = 60$ nm and evolved in the NVT ensemble using the LAMMPS package [3] with implicit solvent (Langevin dynamics) at a temperature of $T = 300$ K. The equations of motion were integrated with a time step of 10 fs for a total simulation time of 5 $\mu$s ($5 \times 10^8$ steps). The radius of gyration ($R_g$) was calculated as

$$R_g^2 = \frac{1}{M_t} \sum_{i=1}^{N} m_i |\mathbf{r}_i - \mathbf{r}_{\rm com}|^2, \qquad \text{(S2)}$$

where $M_t$ is the total mass of the polymer, $\mathbf{r}_{\rm com}$ is the polymer's center of mass, and $m_i$, $\mathbf{r}_i$ are the mass and position of the $i$-th bead, respectively. $R_g$ values were recorded every $10^5$ time steps (1 ns). The first 1 $\mu$s of each trajectory was excluded from the analysis to ensure equilibration. The average radius of gyration obtained was $R_g = 3.9 \pm 0.9$ nm for wild-type NANOG and $R_g = 4.8 \pm 1.2$ nm for the W10A mutant.

#### Cluster simulations of proteins

Simulations of proteins were carried out using the Mpipi coarse-grained model. The simulation proto-

| Region | Length | Residue sequence |
|---|---|---|
| ND | 95 | MSVGLPGPHSLPSSEEASNSGNASSMPAVFHPENYSCLQGSATEMLCTEAASPRPSSEDLPLQGSPDSSTSPKQKLSSPEADKGPEEEENKVLAR |
| HD | 60 | KQKMRTVFSQAQLCALKDRFQKQKYLSLQQMQELSSILNLSYKQVKTWFQNQRMKCKRWQ |
| CD1 | 42 | KNQWLKTSNGLIQKGSAPVEYPSIHCSYPQGYLVNASGSLSM |
| WR | 50 | WGSQTWTNPTWSSQTWTNPTWNNQTWTNPTWSSQAWTAQSWNGQPWNAAP |
| CD2 | 58 | LHNFGEDFLQPYVQLQQNFSASDLEVNLEATRESHAHFSTPQALELFLNYSVTPPGEI |

Table S1. **NANOG sequence across distinct regions.** Amino acid sequences corresponding to the N-terminal domain (ND), homeodomain (HD), C-terminal domain 1 (CD1), tryptophan repeat (WR), and C-terminal domain 2 (CD2) are shown. The second column reports the number of amino acids in each region.

col was adapted from the direct-coexistence approach, a standard method for probing phase separation in biomolecular systems. In this setup, protein molecules are placed within an elongated simulation box that allows for the potential coexistence of dense and dilute phases, if present.

The simulation box was defined as a rectangular prism with its longest axis aligned along the $z$-direction. Periodic boundary conditions were applied in all three dimensions. To minimize finite-size effects and prevent artificial interactions between periodic images, the shorter box dimensions ($L_x$ and $L_y$) were set to at least two to three times the radius of gyration ($R_g$) of a single protein under dilute conditions. The box length $L_z$ was then adjusted to achieve an overall biomolecular concentration corresponding to an average density of approximately 0.1 g/cm$^3$.

We simulated $M = 50$ protein molecules in total. For wild-type NANOG, the simulation box dimensions were $L_x = L_y = 15$ nm and $L_z = 70$ nm, whereas for the W10A mutant the box size was $L_x = L_y = 18$ nm and $L_z = 53$ nm. Each system was first equilibrated at a high temperature ($T = 473$ K) for $5 \times 10^6$ timesteps (approximately 0.5 $\mu$s) to prevent any attractive interactions between polymers and generate a uniform density distribution in the box. Following equilibration, the temperature was quenched to $T = 300$ K, and production simulations were performed in the NVT ensemble using a Langevin thermostat with a relaxation time of 5 ps and an integration timestep of 10 fs, for a total duration of $\sim 2 \times 10^9$ timesteps (2 $\mu$s) for NANOG and $1 \times 10^9$ timesteps for W10A. For each case we ran three independent replicas.

### Data analysis

To assess whether the system had reached equilibrium, we first monitored the total number of intermolecular contacts ($N_{\text{inter,c}}$). Contacts were defined between residues from different proteins when their distance satisfied $r_{ij} \leq 14.5$ Å, corresponding to the shortest cutoff distance of the Wang-Frenkel potential for non-charged particles in the Mpipi model [1]. The time evolution of $N_{\text{inter,c}}$ is shown in Fig. S1(a). Based on this analysis, the initial 16 $\mu$s of the NANOG simulation and the first 7.5 $\mu$s of the W10A simulation were excluded from subsequent analysis.

After equilibration, the simulations revealed that NANOG molecules spontaneously assemble into self-limited clusters, each comprising a well-defined maximum size of approximately 30 proteins. A representative snapshot of the equilibrated system (inset of Fig. S1(a)) shows the formation of two stable clusters (see also Fig. **2**(a) of the main text).

To characterize the internal organization of these assemblies, we computed the intermolecular contact map (Fig. S1(b), inset). Regions of high contact probability, shown in light colors, are concentrated within the WR segment (residues 195–268). A distinct band of interactions is also evident between the WR and the HD (residues 142–159), indicating frequent cross-domain associations. From the WR–WR interactions, we further obtained the distribution of intermolecular connections, defined as the number of NANOG molecules with which a given molecule forms WR-mediated contacts. This distribution follows a Gaussian profile, with an average of seven intermolecular contacts per NANOG molecule within individual clusters (Fig. S1(b)).

At a given timestep, the connectivity between molecules can be represented as a network diagram (see inset of Fig. S1(a)), where each vertex corresponds to a NANOG molecule and edges connect pairs of molecules that form WR-mediated contacts. A connected component is defined as a set of vertices in which any two molecules are linked by one or more connecting paths. Analysis of this network revealed that the system consistently organizes into two main connected components, corresponding to the two clusters observed in the simulations. Occasionally, a few molecules transiently detach from these clusters, forming isolated monomers or dimers. This behaviour was quantified by monitoring the time evolution of the number of molecules in each connected component (see Fig. **2**(c) in the main text). Notably, across all independent replicas, the size of the largest connected component never exceeded 30 molecules, confirming that NANOG forms self-limited clusters.

To examine this self-limiting behaviour and the stability of clusters, we performed a controlled test in which the two NANOG clusters were driven to collide by applying external forces. After the clusters merged, the forces were removed and the system was allowed to relax for 20 $\mu$s.

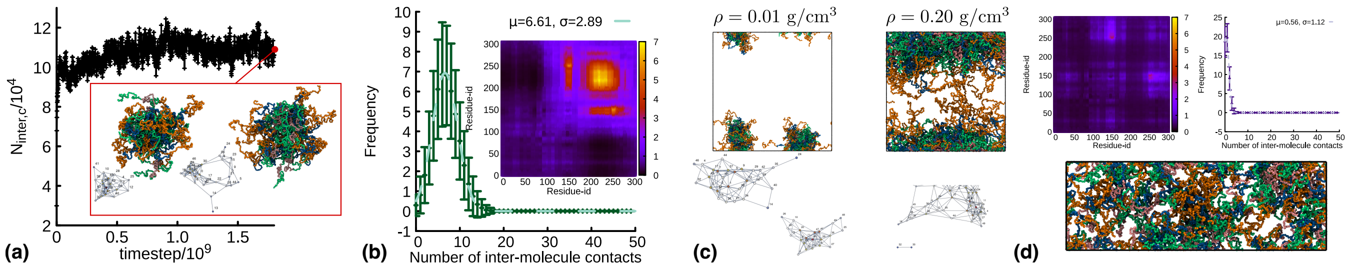


Figure S1. **NANOG forms self-limited clusters**. **(a)** Time evolution of the total number of intermolecule contacts. Inset shows a snapshot from equilibrated simulations of NANOG ($M = 50$, $N = 305$, $T = 300$ K, $\rho = 0.1$ g/cm$^3$). Network diagrams show the molecules in each of the two clusters (29 and 21 molecules, respectively). **(b)** Main: distribution of the number of intermolecular contacts. Inset: residue-level intermolecular contact map. **(c)** Equilibrium configurations from simulations and the corresponding network diagrams. Left: $\rho = 0.01$ g/cm$^3$, showing two clusters containing 29 and 21 molecules, respectively. Right: $\rho = 0.2$ g/cm$^3$. **(d)** Data from simulations of the W10A mutant ($M = 50$, $N = 305$, $T = 300$ K, $\rho = 0.1$ g/cm$^3$), comprising a representative snapshot, the distribution of intermolecular contacts, and the residue-level contact map.

Remarkably, the system spontaneously reorganized into two clusters again, with proteins from the initial aggregates redistributed between the new ones (Fig. **2**(e) of the main text).

In classical phase separation, systems prepared within the coexistence region spontaneously separate into coexisting high- and low-density phases. Although the densities of these phases remain constant, the number of molecules in each phase, and consequently the size of the condensed domains, typically varies with the overall concentration. This naturally raises the question of whether the cluster size observed here depends on box geometry or initial concentration. As shown in Fig. S1(c, left), this is not the case: a system initialized at tenfold lower density ($\rho = 0.01$ g/cm$^3$) in a cubic box still formed two clusters, supporting the self-limited nature of the assembly. At higher densities ($\rho = 0.2$ g/cm$^3$; Fig. S1(c, right)), however, the system transitioned from discrete clusters to a percolating network in which a single aggregate spanned the entire box, marking the onset of gel-like behaviour, consistent with the sample shown in Fig. **1**(c) of the main text.

Finally, we performed simulations of the W10A mutant. Substituting the tryptophan residues with alanine in the WR region resulted in a uniform distribution of molecules throughout the simulation box (Fig. S1(d)). No evidence of cluster formation was observed, underscoring the essential role of the tryptophan residues in mediating NANOG self-association, consistent with previous experimental observations [4].

### III. CRYOEM

3 $\mu$l of purified recombinant NANOG (see next section for purification method) at a concentration of 1 mg/ml were spotted on Quantifoil 2/2 grids with blotting for 3.5 seconds and blotted on a Vitrobot instrument with a blot force of 4. 6407 multi-frame movies in total were collected on a JEOL CRYOARM300 microscope equipped with a DE64 detector. The data collection was performed using Serial EM in linear mode using a pixel size of 1 Å/pixel and 58 e-/Å$^2$. The movies were motion corrected using MotionCorr and the CTF estimation was carried out using the CTFFIND 4.1 function in RELION 5.0 [5]. 1,726 movies were selected based on motion correction and CTF parameters. 1,036 particles were picked from the micrographs manually, extracted using a box size of 384x384 pixel and coarsened to 64x64 pixels prior to 2D classification. These class averages were then used as references for referenced autopicking of the data set producing a particle set of 1,048,156 particles. Following extraction, classification and particle selection in Relion 5.0, a final dataset of 121,110 particles was classified into 50 classes using a 240 Å  mask.

### IV. PURIFICATION OF RECOMBINANT NANOG

Full-length Nanog, W10A and N51A were expressed in pET15b and purified according to established procedures described in Refs. [4] and  [6]. We briefly summarize the methods here.

#### Purification of recombinant NANOG

BL21(DE3) bacteria were transformed with NANOG-carrying gene under the control of an inducible promoter. The over-expression was triggered by 1mM isopropyl $\beta$-d-1-thiogalactopyranoside (IPTG) and grown for 3 hours at 37°C. See Ref. [6] for more details.

The protein was then purified in the presence of urea using nickel resin columns. Purified protein was concentrated in Vivaspin devices as in Ref. [6]. The protein was kept in denaturing buffer (25mM Hepes pH7.6, 150mM NaCl, 7M urea, 10mM imidazole, 10mM beta-

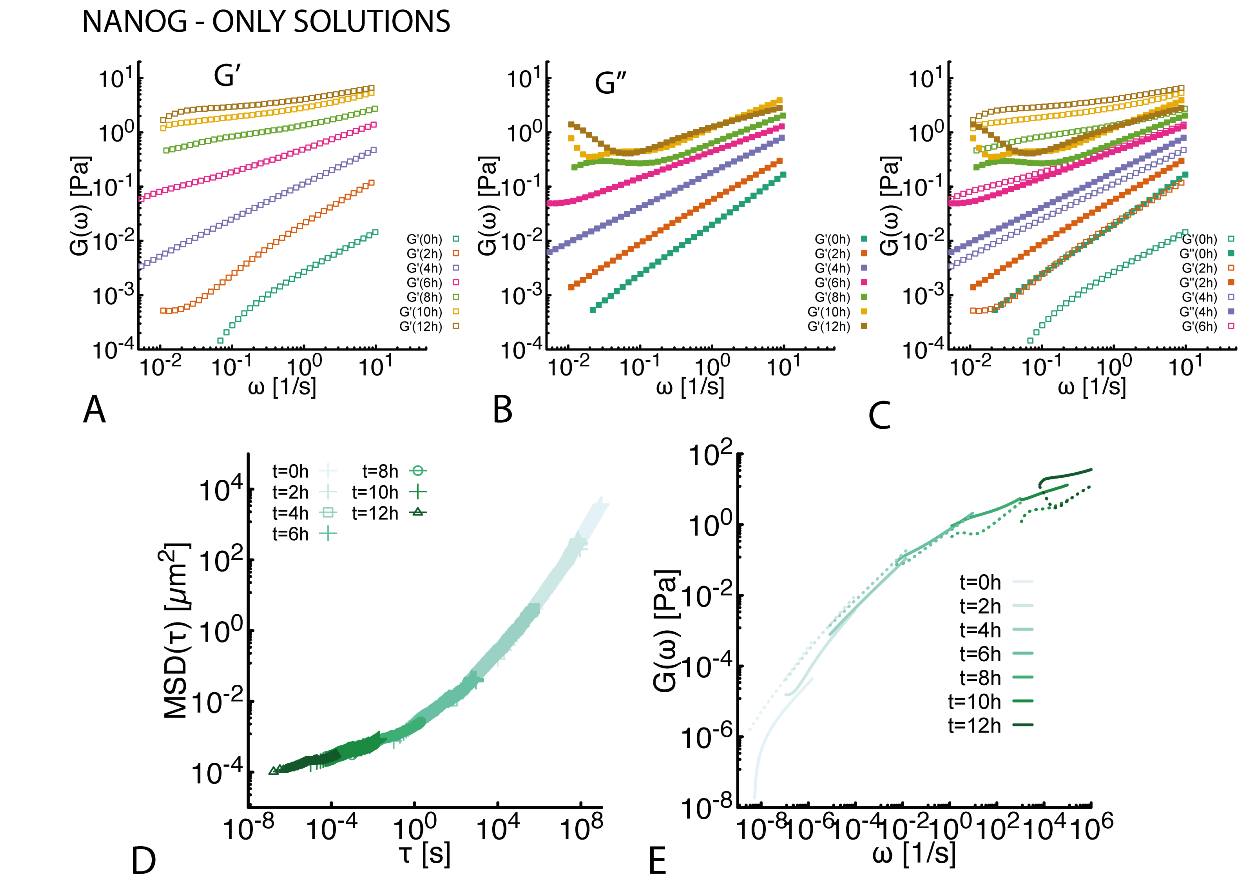


Figure S2. **Additional rheology curves for NANOG WT only solution during aging**. **A** All $G'$ and **B** $G''$ curves obtained from tracers' MSD during aging of NANOG WT solutions at 1mM incubated at 37C for 12h. **C** Plot reporting $G'$ and $G''$ curves together. **D-E** Time-cure superpositions of MSDs and shear moduli of NANOG WT only solution at 1mM during aging at 37C.

mercaptoethanol). The protein was then run on an SDS-PAGE gel to confirm its purity and its concentration was measured by Bradford assay.

Concentrated protein was refolded by diluting denatured protein (10 mg/ml) in refolding buffer (10mM Hepes pH7.6, 50mM KCl, 10mM NaCl, 0.4mM EDTA, 10% glycerol) 1 in 10 (final 1 mg/ml) and incubating at room temperature for 30 minutes. Protein was stored refolded at 4°C for up to 2 weeks and diluted in refolding buffer as needed.

#### Electromobility shift assay (EMSA)

The DNA probe used was a IRDye 680nm labelled 26bp sequence from the Tcf3 promoter [7] (`TAAAC CTGTTAATGGGAGCGCATTG`). Final protein concentrations were as indicated in the figure. Samples were incubated for 30 min at room temp and analyzed on 1.5% agarose TBE gels. The gel was imaged at 700nm.

### ADDITIONAL MICRORHEOLOGY EXPERIMENTS AND ANALYSIS

#### Time-cure superposition during aging of NANOG solutions

In Fig. S2 we report the time-cure superposition curves for the MSDs and $G'$ and $G''$ for solutions of WT NANOG at 1mM at 37°C. In the time-cure superposition plots for the MSDs, each curve at different "curing time" (aging time $t_a$) is shifted by a lagtime $\tau_c(t_a)$ and MSD value $m_c(t_a)$. The corresponding shear moduli are shifted in frequency by $\omega_c(t_a) = 1/\tau_c(t_a)$ and by a value $G_c(t_a)$.

#### Time-cure superposition during aging of DNA-NANOG solutions

In Fig. S3 we report the time-cure superposition curves for the MSDs and $G'$ and $G''$ for solutions of $\lambda$DNA mixed with NANOG. DNA concentration is 7.9nM while NANOG concentration is 7.9$\mu$M. The solution is kept at 37°C overnight and movies of tracers are recorded every 2 hours. In the time-cure superposition plots for the MSDs, each curve at different "curing time" (aging time $t_a$) is shifted by a lagtime $\tau_c(t_a)$ and MSD value $m_c(t_a)$. The corresponding shear moduli are shifted in frequency by $\omega_c(t_a) = 1/\tau_c(t_a)$ and by a value $G_c(t_a)$. The superposed curves do not follow a simple Maxwell fluid model as in Ref. [8] as they display multiple relaxation times where $G'$ and $G''$ crossover. This is consistent with the NANOG-DNA solutions displaying strong gel-like behaviour which was not observed in Ref. [8]. We hope to further analyse this behaviour in the future.

### Gelation of DNA-NANOG solutions require long DNA

In figure S4 we show MSDs of tracers embedded in a solution of $\lambda$DNA cut by HaeIII. This restriction enzyme cleaves DNA in 130 fragments with an average length of 1 kbp. When we incubate the samples with 7.9nM of HaeIII-cut $\lambda$DNA and 7.9$\mu$M NANOG WT at room temperature we do not observe gelation. This evidence suggests that the gelation is caused by NANOG micelles stabilizing crosslinks between long DNA molecules. In the case of short, not entangled DNA, NANOG micelles bind DNA fragments and form freely diffusing particles.

### Behaviour of NANOG and DNA-NANOG solutions at Room Temperature

In Fig. S5a, we show that NANOG-only solutions prepared at 1 mM and incubated for 12 h do not exhibit a significant onset of elasticity; rather, they display a substantial increase in viscosity over the course of the incubation period. The increase in viscosity suggests that the NANOG particles are increasing in hydrodynamic size but do not form a percolating network of micelles.

In Fig. S5b, we show that DNA-NANOG solutions prepared at 1:1000 DNA:protein ratio (7.9 nM DNA, 7.9 $\mu$M NANOG WT) and incubated at room temperature for 12h do show gelation. We interpret this behaviour as in line with the data reported in the main text, NANOG forms micelles that bind DNA and crosslink entangled DNA.

### Validation that microrheology is independent of tracer size and coatings

In Fig. S6 we compare the MSDs of tracers in NANOG-only and DNA-NANOG solutions past the gel point. We consider three types of tracers with different sizes and coatings. Specifically, 1 $\mu$m Fluoresbrite BB Carboxylate Microspheres (Polyscience), passivated by resuspending them in high concentration BSA solution followed by washing before use; (ii) 2 $\mu$m PEG-coated (MW 5k) Fluoresbrite BB Carboxylate Microspheres (Polyscience); (iii) 3 $\mu$m Latex beads, polystyrene from Sigma. In the plot one can appreciate that once rescaled by their size, the MSDs collapse on top of each other, indicating that there is no size or coating dependence, i.e. our results are independent of tracer choice for the sizes and coatings considered.

* Joint first author

[1] J. A. Joseph, A. Reinhardt, A. Aguirre, P. Y. Chew, K. O. Russell, J. R. Espinosa, A. Garaizar, and R. Collepardo-Guevara, Nature Computational Science **1**, 732 (2021).
[2] I. Chambers, D. Colby, M. Robertson, J. Nichols, S. Lee, S. Tweedie, and A. Smith, Cell **113**, 643 (2003).
[3] A. P. Thompson, H. M. Aktulga, R. Berger, D. S. Bolintineanu, W. M. Brown, P. S. Crozier, P. J. in 't Veld, A. Kohlmeyer, S. G. Moore, T. D. Nguyen, R. Shan, M. J. Stevens, J. Tranchida, C. Trott, and S. J. Plimpton, Comp. Phys. Comm. **271**, 108171 (2022).
[4] N. Mullin, A. Yates, A. Rowe, B. Nijmeijer, D. Colby, P. Barlow, M. Walkinshaw, and I. Chambers, Biochemical Journal **411**, 227 (2008).
[5] A. Burt, B. Toader, R. Warshamanage, A. von Kügelgen, E. Pyle, J. Zivanov, D. Kimanius, T. A. M. Bharat, and S. H. W. Scheres, FEBS Open Bio **14**, 1788 (2024).
[6] N. P. Mullin, A. Gagliardi, L. T. P. Khoa, D. Colby, E. Hall-Ponsele, A. J. Rowe, and I. Chambers, Journal of Molecular Biology **429**, 1544 (2017).
[7] R. Jauch, C. K. L. Ng, K. S. Saikatendu, R. C. Stevens, and P. R. Kolatkar, Journal of Molecular Biology **376**, 758 (2008).
[8] L. Jawerth, E. Fischer-Friedrich, S. Saha, J. Wang, T. Franzmann, X. Zhang, J. Sachweh, M. Ruer, M. Ijavi, S. Saha, J. Mahamid, A. A. Hyman, and F. Jülicher, Science **370**, 1317 (2020).

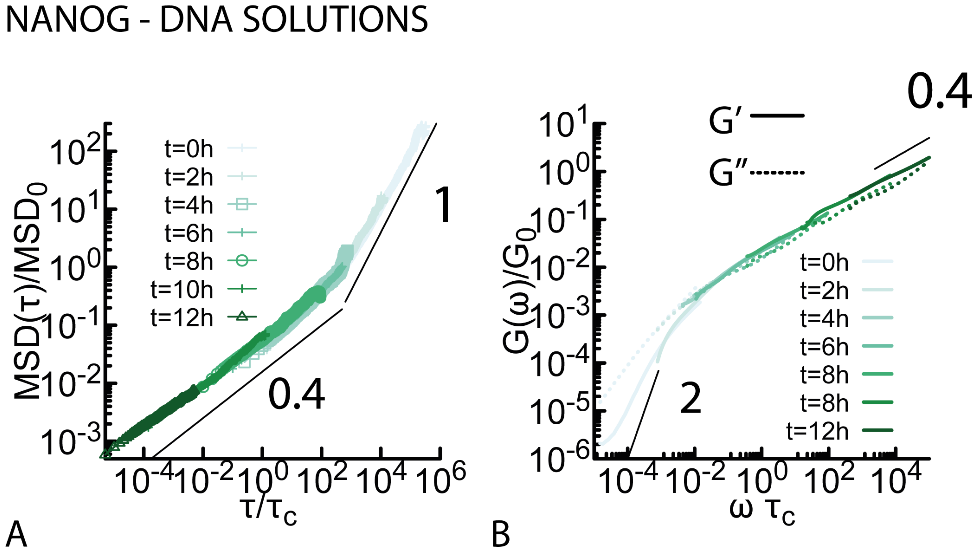


Figure S3. **Rheology analysis of NANOG-DNA mixtures**. **A-B** Time-cure superposition of MSDs and shear moduli of NANOG-DNA mixtures during aging. DNA concentration 7.9 nM, NANOG WT concentration 7.9 $\mu$M, incubated at 37°C for 12h.

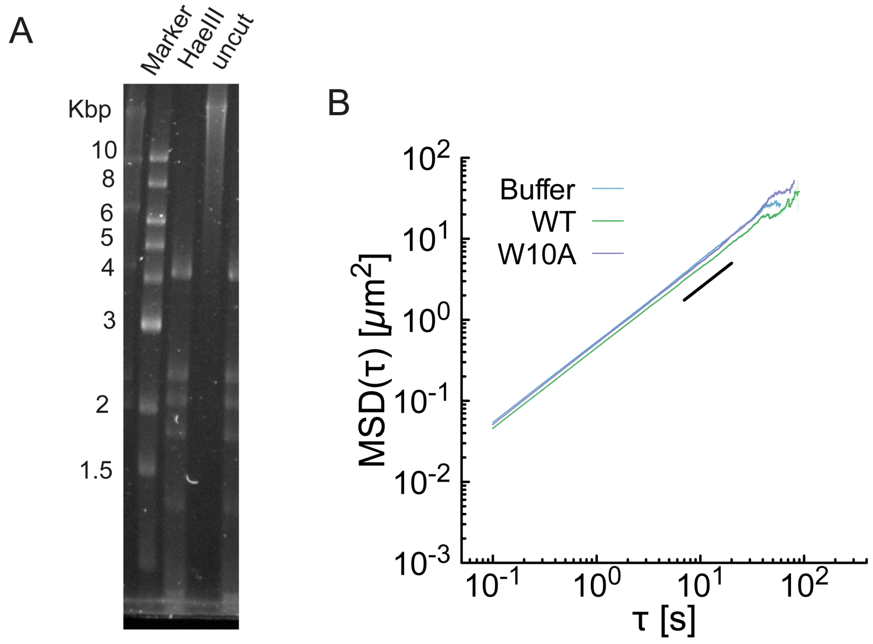


Figure S4. **Gelation of DNA-NANOG solutions require long DNA molecules**. **A** Gel electrophoresis of $\lambda$DNA and $\lambda$DNA cut with HaeIII. **B** MSD of tracers embedded in a buffer solution and compared with a solution of 7.9 nM HaeIII-cut $\lambda$DNA and NANOG WT at 7.9 $\mu$M at room temperature. In these conditions we do not observe gelation.

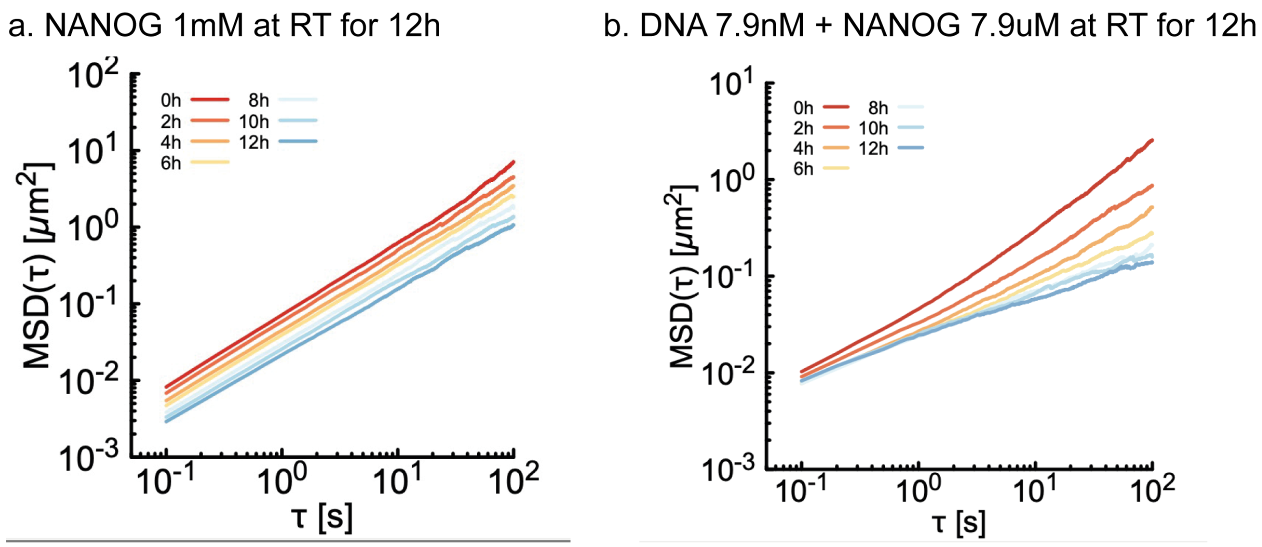


Figure S5. **NANOG and NANOG-DNA solutions aging at room temperature**. **a.** MSD of tracers embedded in a solution of NANOG at 1mM with overnight incubation at room temperature. **b.** MSD of tracers embedded in a DNA:NANOG solution at 7.9nM:7.9$\mu$M (1:1000) ratio. The solution is incubated at room temperature for 12h.

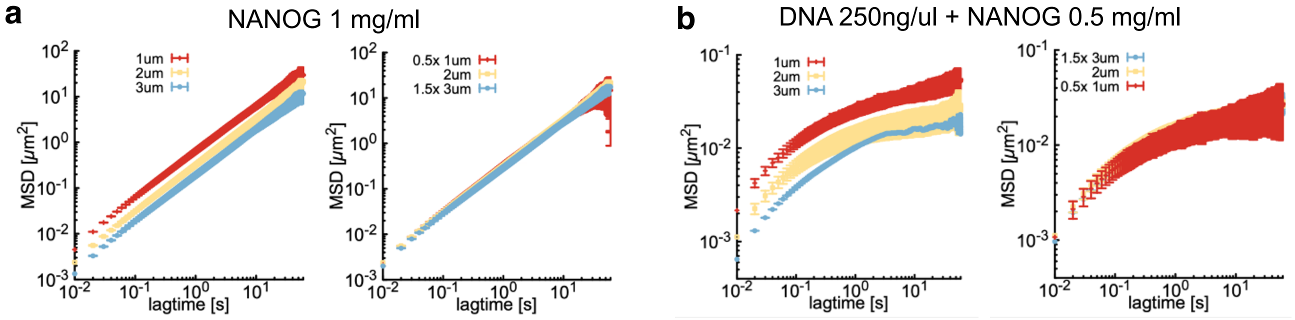


Figure S6. **Bead size and coating do not affect microrheology measurements**. **a, Left** MSD of different tracers embedded in a WT NANOG-only system at 1 mg/ml (28 $\mu$M). **(a, Right)** MSDs rescaled by the size of the tracers relative to the 2 $\mu$m tracer. The collapse of the curves indicate that there is no significant size or coating dependence. **b, Left** MSD of different tracers embedded in a NANOG-DNA system at 250 ng/ul (7.9 nM) $\lambda$DNA and 0.5 mg/ml (7.9 $\mu$M) (WT NANOG) beyond the gel point. **b, Right** MSDs rescaled by the size of the tracers relative to the 2 $\mu$m tracer. The collapse of the curves indicate that there is no significant size or coating dependence. The tracers in these experiments are as follows: (i) 1 $\mu$m Fluoresbrite BB Carboxylate Microspheres (Polyscience), passivated by resuspending in high concentration BSA solution followed by washing before use; (ii) 2 $\mu$m PEG-coated (MW 5k) Fluoresbrite BB Carboxylate Microspheres (Polyscience); 3 $\mu$m Latex beads, polystyrene from Sigma.
